# Mapping Functional Homologies Between Human and Marmoset Brain Networks Using Movie-Driven Ultra-High Field fMRI

**DOI:** 10.1101/2025.05.07.652702

**Authors:** A. Zanini, A. Dureux, R.S. Menon, S. Everling

## Abstract

Naturalistic stimuli, such as movies, offer a powerful tool for probing functional brain organization across species. Using movie-driven functional magnetic resonance imaging (md-fMRI), we recorded brain activity in humans and awake marmosets exposed to the same dynamic audiovisual stimulus. We applied tensor independent component analysis (tICA) to identify functional networks in each species, hierarchically clustered these components, and examined their within- and between-species temporal correlations to assess functional homologies. We found strong interspecies correspondence in core sensory networks, particularly those involved in visual and auditory processing, suggesting conserved mechanisms for sensory integration. In contrast, networks associated with higher-order cognition, including prefrontal and temporoparietal areas, were observed primarily in humans, highlighting species-specific specializations. These findings demonstrate the value of naturalistic paradigms and data-driven approaches in revealing both shared and divergent brain network architectures. By openly sharing our data and analysis pipelines, we aim to support future comparative studies and advance the marmoset as a model for investigating the evolutionary foundations of brain function.

## Introduction

The common marmoset (*Callithrix jacchus*) is gaining traction as a primate model in neuroscience due to its small size, rapid maturation, and rich cognitive and social behaviors (1–3). The New World marmoset shares key features with Old World monkeys and humans, including a granular prefrontal cortex (4), a complex visual system (5), and comparable visuomotor behaviors (6,7). Recent studies have also uncovered an extensive network dedicated to processing conspecific vocalizations, consistent with the species’ reliance on social communication (8–10).

As interest in the marmoset grows, understanding the functional organization of its brain — and how it compares to the human brain — becomes increasingly important. Functional magnetic resonance imaging (fMRI) provides a powerful tool for this purpose, enabling whole-brain recordings during task-based or resting-state paradigms. Prior studies have successfully used both approaches to compare human and marmoset brain function, revealing homologous resting-state networks (11–14) and activations in response to shared stimuli (15,16).

However, both fMRI approaches have limitations. Resting-state fMRI can reveal functional connectivity across brain regions (17,18) but lacks the specificity to identify networks driven by behaviorally relevant stimuli. Moreover, although resting-state activity is spontaneous, it is also highly dependent on the subject’s internal state — which is often unknown or uncontrolled — making interspecies comparisons of functional networks difficult. Task-based fMRI, by contrast, allows for direct comparison of stimulus-driven responses and has been used extensively in humans (16,19), macaques (20,21), and marmosets (9,22–26). Yet this method is limited by the impracticality of designing and repeating numerous functional localizers across the entire brain, especially in non-human primates.

Movie-driven fMRI (md-fMRI) has emerged as a promising alternative. Naturalistic movies provide dynamic audiovisual input that elicits robust and reliable neural responses across sensory and cognitive systems in awake animals. Crucially, such stimuli maintain the engagement of non-human primates during long fMRI sessions and offer a more ecologically valid alternative to traditional paradigms. Studies in humans (27–31), macaques (32,33), and marmosets (14) have shown that movies can reliably drive widespread, selective brain activity. In this study, we presented a naturalistic movie featuring diverse visual and auditory stimuli to eight marmosets and nineteen humans. Using tensor independent component analysis (tICA), a data-driven approach well suited to naturalistic paradigms, we identified 20 functional components in each species. We then examined their temporal dynamics within and across species to identify potential functional homologues. Our findings reveal both conserved and divergent patterns of brain activity, shedding light on the evolutionary organization of sensory and cognitive networks in primates.

## Methods

### Common marmosets

All procedures followed Canadian Council on Animal Care guidelines and were approved by the University of Western Ontario Animal Care Committee. Eight adult common marmosets (4 females; age: 28–45 months, mean: 39.8 months) participated in the study. Each animal underwent surgery to implant a PEEK head post (34), following the protocol detailed in our recent methods article (35). Briefly, under gas anesthesia (0.5–3% isoflurane), the skull was exposed, prepared with adhesive resin (All-Bond Universal, Bisco), and affixed with a resin composite (Core-Flo DC Lite, Bisco). Vital signs were continuously monitored throughout surgery. After two weeks of recovery, animals were acclimated to the head-fixation system over three weeks in a mock MRI environment.

### Human participants

Nineteen healthy, self-reported right-handed participants (11 females; age: 25–45 years, mean: 32.7 years), with normal or corrected-to-normal vision and no neurological or psychiatric history, were recruited. Fourteen had previous fMRI experience. All participants provided written informed consent, and the study was approved by the University of Western Ontario Human Research Ethics Board.

### Stimuli

Marmosets and humans viewed a 33-minute naturalistic movie composed of alternating baseline periods (5:50 min total) and excerpts from two nature documentaries: *Monkey Kingdom* (Disneynature, Spanish narration) and *Hidden Kingdoms – Urban Jungles* (BBC, English narration). The movie contained a wide variety of visual (e.g., marmosets, humans, animals, cityscapes, landscapes) and auditory stimuli (e.g., speech, vocalizations, music, environmental sounds). Baseline periods displayed a fixation target (black circle, 0.36° visual angle) on a gray screen without audio.

### Experimental Setup

#### Marmosets

During scanning, awake animals were seated in a custom 3D-printed sphinx-style chair with head fixation using the headpost (34,35). Visual stimuli were rear-projected onto a screen 119 cm from the eyes using a Sony VLP-FE40 LCSD projector reflected off a front-surface mirror and were presented using PowerPoint, synchronized with the MRI TTL pulse via a Raspberry Pi (model 3B+) running custom Python software. Auditory stimuli were delivered via Sensimetrics S14 tubes, secured with earplugs and veterinary bandage. An MRI-compatible camera (MRC Systems) monitored the animals, though eye tracking was unreliable due to partially closed eyelids. A drop of marshmallow-flavored liquid reward was delivered every 4.5 s via a tube to maintain alertness. Each marmoset watched the movie five times across five sessions.

#### Humans

Participants lay supine in the scanner and viewed the stimulus through a mirror mounted on the head coil. Visual stimuli, projected using an Avotech SV-6011 projection system, were presented using PowerPoint, synchronized with the MRI TTL pulse via a Raspberry Pi (model 3B+) running custom Python software. Audio was delivered through Sensimetrics T14 tubes, with participants confirming acceptable volume prior to scanning. Each subject viewed the movie once.

### MRI data acquisition

#### Marmosets

Imaging was performed at 9.4T (31-cm bore Varian magnet interfaced to a Bruker Avance NEO console) with a custom 15-cm gradient coil and eight-channel receive coil inside a quadrature birdcage transmit coil. Functional images were acquired in five sessions per animal using gradient-echo EPI (TR = 1.5 s, TE = 15 ms, flip angle = 40°, FOV = 64 × 48 mm, matrix = 96 × 128, resolution = 0.5 mm isotropic, 42 axial slices, bandwidth = 400 kHz, GRAPPA = 2). Additional EPI runs with reversed phase encoding were collected for distortion correction. A T2-weighted structural scan was acquired in one session (TR = 7 s, TE = 52 ms, FOV = 51.2 x 51.2 mm, bandwidth 50 kHz, resolution = 0.133 × 0.133 × 0.5 mm).

#### Humans

Data were acquired at 7T (Siemens Magnetom MRI Plus) with a 32-channel receive and 8-channel parallel transmit coil. Functional images were acquired using multi-band EPI (TR = 1.5 s, TE = 20 ms, flip angle = 30°, FOV = 208 × 208 mm, matrix = 104 × 104, resolution = 2 mm isotropic, 62 slices, GRAPPA = 3, multi-band factor = 2). Field maps were derived from magnitude and dual-phase images. MP2RAGE structural images were collected (TR = 6 s, TE = 2.13 ms, TI1/TI2 = 800/2700 ms, resolution = 0.75 mm isotropic).

### fMRI data preprocessing

#### Marmoset Data

Preprocessing was performed using AFNI (36) and FSL (37). DICOMs were converted with dcm2niix, reoriented (fslswapdim, fslorient), and distortion-corrected (topup, applytopup). Volumes were despiked, slice-time corrected, motion-corrected, and spatially smoothed (1.5 mm FWHM). Nuisance regression included motion parameters. Temporal filtering (0.01–0.08 Hz) was applied using 3dBandpass. Functional data were registered to individual anatomy (FLIRT), then to the NIH marmoset template (38) using ANTs (39).

#### Human Data

Preprocessing was done with SPM12 (Wellcome Center for Human Neuroimaging, London, UK) and AFNI. After conversion with dcm2niix, images were realigned, slice-time corrected, field map–corrected, coregistered to structural MP2RAGE, normalized to MNI space, and smoothed (6 mm FWHM). Temporal filtering (0.01–0.08 Hz) matched the marmoset pipeline.

### Statistical Analysis

We applied tICA using MELODIC (40) with a 20-component cutoff for each species, using the entire 33-minute movie. This method decomposes data into spatially and temporally independent components, enabling identification of stimulus-driven functional networks without relying on predefined templates. Twelve components in marmosets and fourteen in humans were classified as non-noise based on spatial and spectral criteria.

We computed within- and between-species correlation matrices of component timecourses and performed hierarchical clustering (squared Euclidean distance, Ward’s method) in R (41) using the cluster (42), dendextend (43), and ggdendro (44) packages. These analyses allowed us to explore the organization of functional networks and to investigate interspecies relationships. This approach is displayed in Figure 1.

**Figure 1.**
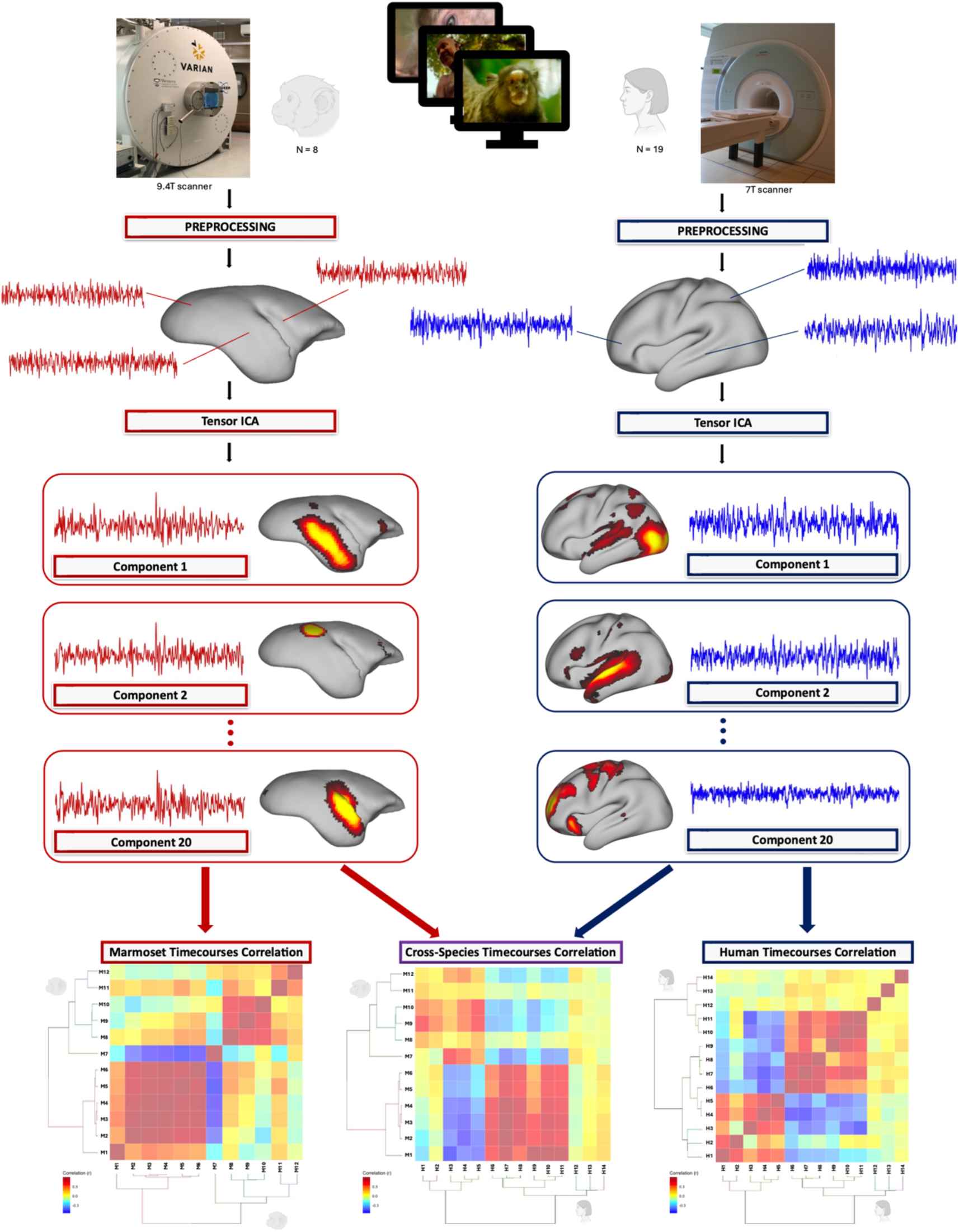
Experimental design and stimulus structure. Subjects viewed a 33-minute movie composed of alternating baseline and movie blocks. Baseline periods (totaling 5:50 minutes) consisted of a fixation point on a gray background with no audio. Movie segments included clips from two nature documentaries, featuring a variety of species and environments. The film contained naturalistic visual and auditory stimuli, including conspecific and heterospecific vocalizations, human speech, music, and environmental sounds. At the top, the 7 Tesla and 9.4 Tesla scanners used for fMRI sessions in humans (n = 19) and common marmosets (n = 8, with 5 functional runs per marmoset) are shown. On the left and right sides of the figure is represented the statistical approach used for both marmosets (left) and human (right). After preprocessing, individual fMRI maps were analyzed using tensor independent component analysis (tICA), extracting 20 independent components per species. The timecourses of non-noise components (n = 12 for marmosets and n = 14 for humans) were then correlated within and across species, generating correlation matrices (example shown at the bottom). Additionally, these functional components were clustered using hierarchical clustering to further explore functional network relationships in both species.

To assess spatial coverage, non-noise components were binarized and averaged to generate probability maps. These were compared with temporal signal-to-noise ratio (tSNR) maps to evaluate recruitment consistency and signal quality.

## Results

### Marmoset tICA Components

Of the 20 components produced by tensor ICA (tICA) decomposition of the marmoset data, 12 were classified as functional (non-noise) based on spatial distribution and temporal power spectra (Figure 2). These components encompassed a broad range of sensory, motor, and associative brain areas, and were grouped into networks based on anatomical distribution and functional relevance.

**Figure 2.**
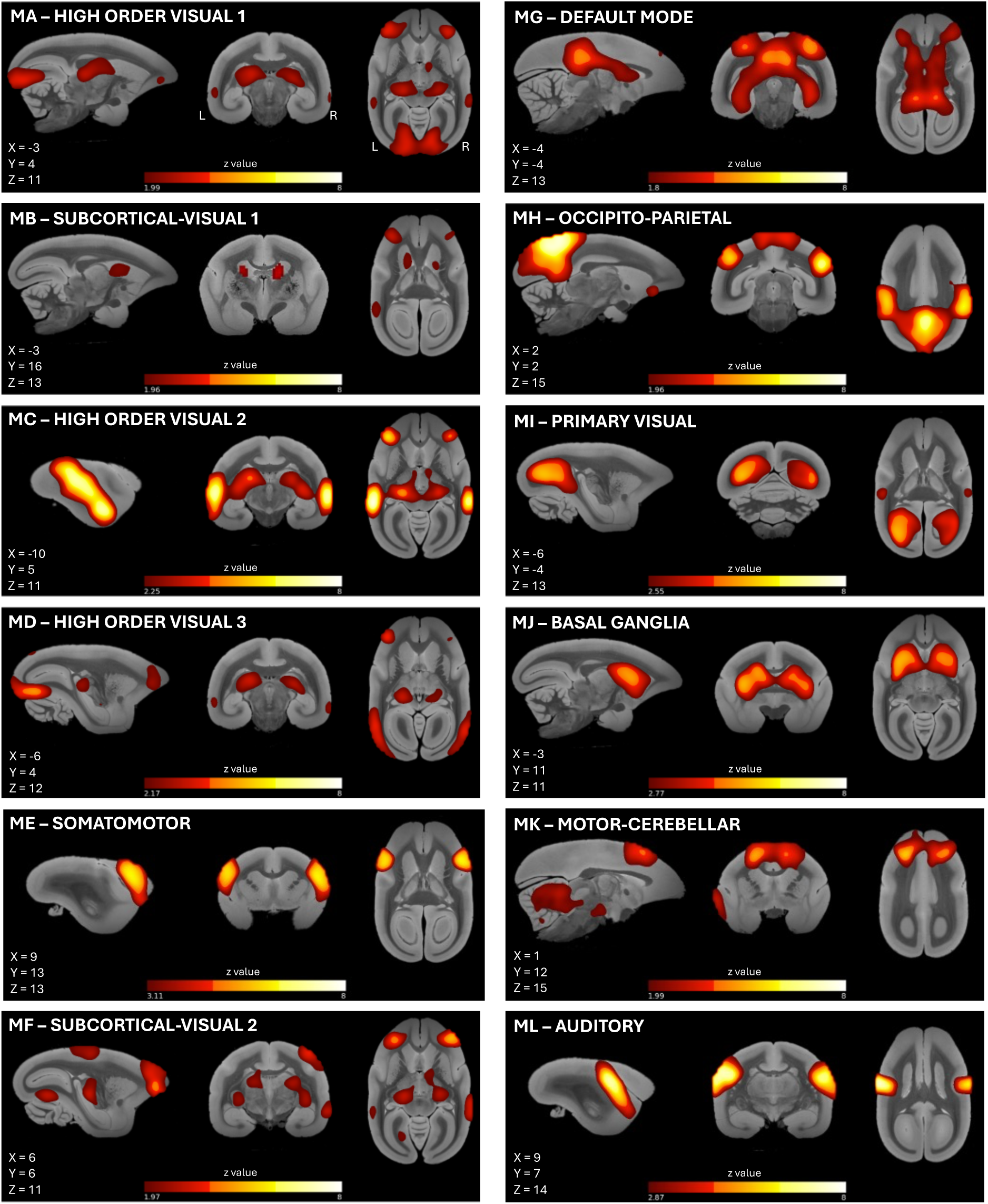
Spatial maps of the 12 functional marmoset tICA components. Components are represented on sagittal, coronal, and transversal slices of a high-resolution template of the marmoset brain (38). Brain coordinates of the presented perspective are reported in the left bottom corner of each panel. Components include higher-order visual (HOV), subcortical-visual (SUV), somatomotor (SOM), default mode (DMN), occipitoparietal (OP), primary visual (PVIS), basal ganglia (BG), motor-cerebellar (MCE), and auditory (AUD) networks. Components were selected based on spatiotemporal features and visual inspection. Color indicates voxel-wise z-scores, thresholded at p < 0.5 (posterior probability that a voxel belongs to the active distribution rather than noise).

Three components (MA, MC, and MD) were categorized as **higher-order visual networks (HOV)** given their anatomical distribution and alignment with prior resting-state studies (13,45,46). These exhibited strong bilateral activation in occipitotemporal visual areas as well as prefrontal cortex regions, including areas 8Av, 8C, 45, and 47. Subcortical activation in these networks included the pulvinar and superior colliculus, indicating integration of visual and attentional processing. Two additional components (MB and MF) were identified as **subcortical-visual networks (SUV)**. These shared overlap with the HOV components in both visual and prefrontal cortex but extended further into subcortical regions, including the caudate, putamen, and thalamus, and recruited parietal areas such as LIP, AIP, PE, PG, and PGM. This pattern suggests involvement in visuomotor integration and spatial attention.

A single component (ME) was classified as a **somatomotor network (SOM)**, with strong activation in primary somatosensory cortex (areas 1/2, 3a, and 3b), motor cortex (area 4ab), and ventral premotor cortex (areas 6Va and 6Vb). Its spatial pattern closely matches the somatomotor ventral component previously described in the literature (12,13,45,46), likely reflecting widespread engagement of sensorimotor regions during the observation of behaviorally relevant stimuli in the movie. Component MG displayed a pattern consistent with a **default mode network (DMN)** (11,13,45–48), with activation in posterior cingulate, medial parietal areas (MIP, LIP, OPt), and dorsomedial prefrontal cortex (areas 6DC, 6DR, 8aD, and 8C), along with subcortical regions such as the hippocampus and caudate. While structurally reminiscent of the primate DMN, its functional properties differed, as discussed further below.

Component MH extended across medial occipital cortex, orbitofrontal cortex, and the putamen and was labeled an **occipitoparietal network (OP)**, likely involved in integrating visual and contextual information. Component MI was more spatially restricted to early visual areas, including V1 and MT, and was identified as a **primary visual network (PVIS)** (13,45,46). In contrast to these cortical-dominant components, MJ exclusively encompassed subcortical structures—putamen, caudate, globus pallidus, and ventral thalamus—without cortical involvement, defining a **basal ganglia network (BG)** (12,13,45). Component MK, the **motor-cerebellar network (MCE)**, revealed co-activation of cerebellar regions, motor and premotor cortex (4ab, 6DC, 6DR, and 6M), and the periaqueductal gray, possibly reflecting coordinated sensorimotor integration. Lastly, component ML was localized to the core, belt, and parabelt auditory cortex and was labeled the **auditory network (AUD)**, consistent with prior studies using resting-state data in marmosets (12,13,45,46).

Figure 3A illustrates that frequently recruited voxels were most prominent in prefrontal, auditory, and visual cortices, as well as in subcortical structures like the caudate and superior colliculus. These patterns were not simply a byproduct of data quality, as comparison with the temporal signal-to-noise ratio (tSNR) map (Figure 3B) confirmed that component distributions did not merely reflect regions of high SNR.

**Figure 3.**
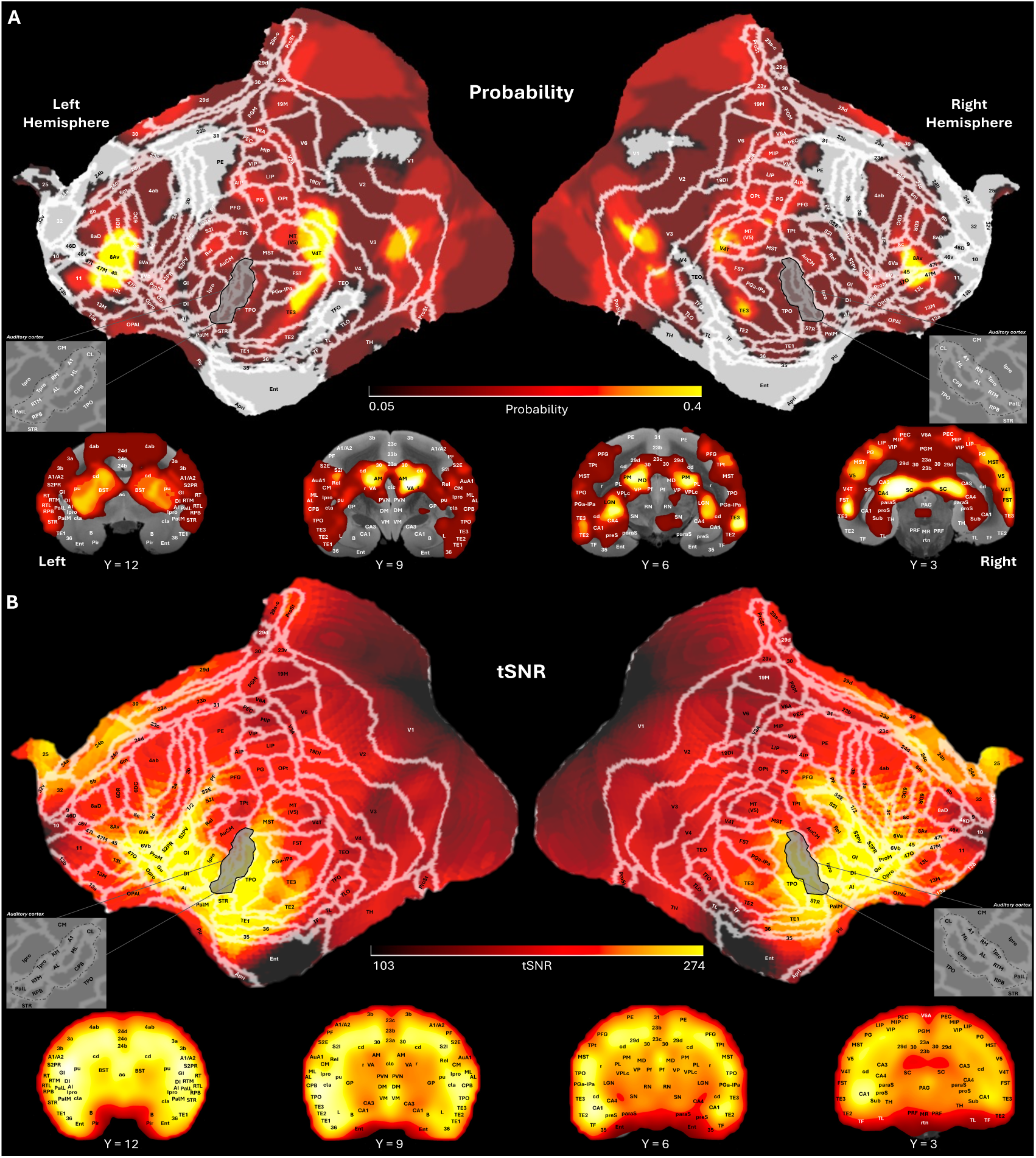
Spatial distribution and tSNR of marmoset components. (A) Probability map showing the number of functional components in which each voxel was active (thresholded as indicated in Figure 2). Voxels most frequently recruited across components are concentrated in prefrontal, auditory, visual, and subcortical regions. (B) Mean temporal signal-to-noise ratio (tSNR) map across marmosets showing the average quality of the BOLD signal across the brain, with higher tSNR values represented in yellow and lower values in black. The similarity between frequently recruited regions and high-tSNR areas confirms that component patterns were not driven solely by signal quality. In both panels, the maps are shown on flat surface representations of the marmoset brain, with white lines delineating the Paxinos parcellation (76) of the NIH marmoset brain atlas (38), and on coronal slices of a marmoset brain anatomical image at different inter-aural levels.

### Marmoset Component Relationships

To assess the interrelationships among the functional networks, we computed temporal correlations between component timecourses and performed hierarchical clustering (Figure 4A). The resulting dendrogram revealed four distinct network groupings, which were mirrored in the within-species correlation matrix (Figure 4B).

**Figure 4.**
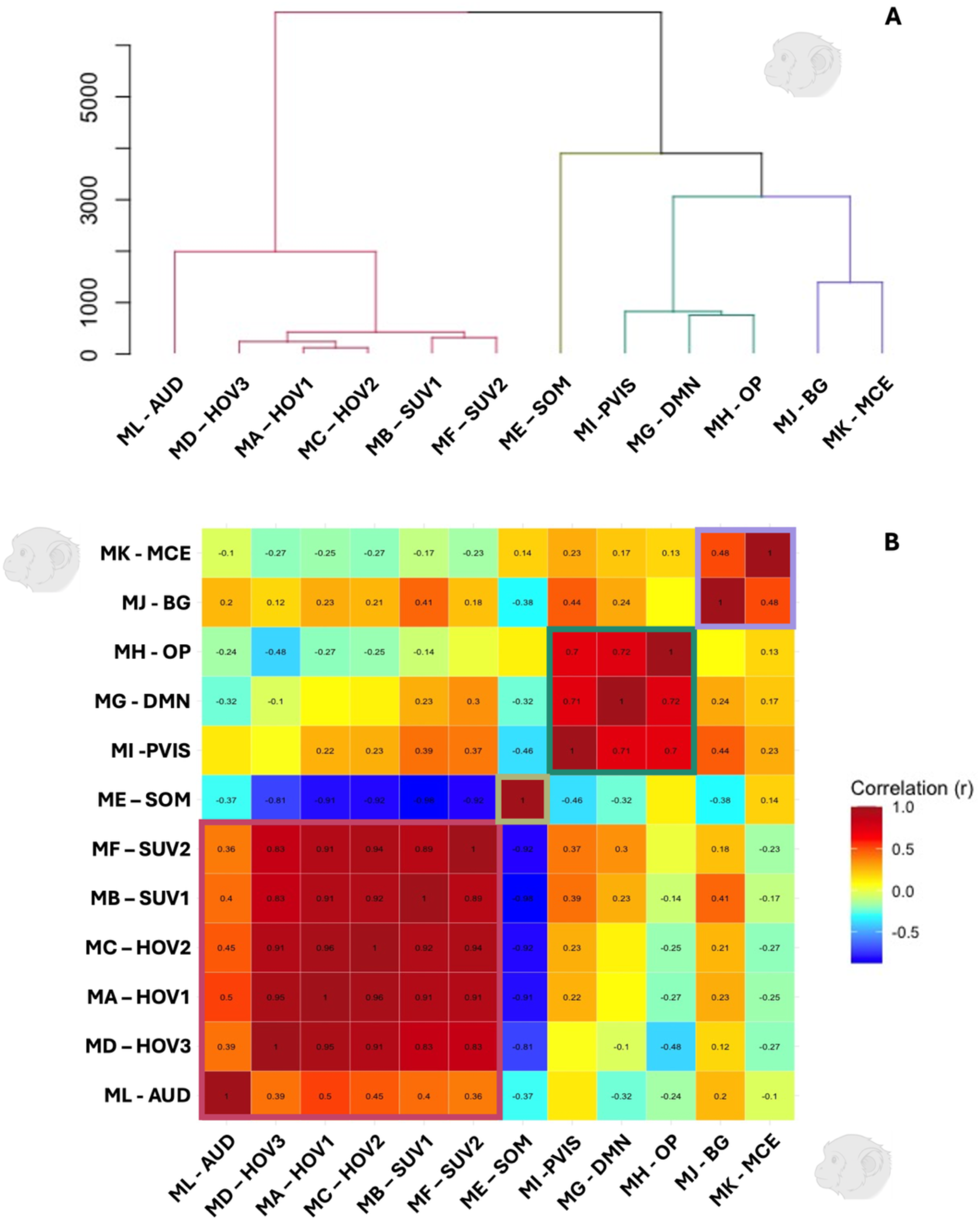
Marmoset component clustering and temporal relationships. (A) Hierarchical clustering dendrogram of the 12 functional marmoset components using squared Euclidean distance and Ward’s linkage. Colors indicate functional clusters: multisensory (red), somatomotor (green), visual-default (turquoise), and subcortical-motor (purple). (B) Within-species correlation matrix showing pairwise temporal correlations (Pearson’s r) between component timecourses. Components are ordered based on the sequence specified by the hierarchical clustering tree. Networks within the same functional group show strong positive correlations, while cross-cluster relationships are more variable, including anticorrelations between somatomotor and multisensory networks. Clusters identified through hierarchical clustering are highlighted on the matrices with colored squares, following the same color scheme presented in panel A. Red shades indicate positive correlations, while blue shades represent negative correlations. Only correlations with an absolute r value greater than 0.10 are displayed.

The first cluster consisted of **multisensory networks**, grouping together the HOV, SUV, and AUD components. HOV and SUV components were very highly inter-correlated (r = 0.83–0.96), suggesting coordinated engagement of visual systems during naturalistic movie viewing. The auditory component (AUD) showed moderate correlations with both HOV and SUV components (r = 0.36–0.50), indicating cross-modal interactions and integrative processing across sensory modalities.

A second, largely independent cluster contained the SOM network, which showed weak correlation with the motor-cerebellar component (MCE; r = 0.14) and strong anticorrelations with all other functional components (r = –0.32 to –0.96). A third group—comprising the visual-default cluster—included the PVIS, DMN, and OP components, which were internally well correlated (r = 0.70–0.72), showed modest correlations with subcortical-visual components, and anticorrelated with the auditory network. Their correlations with the higher-order visual networks were more heterogeneous: PVIS and DMN showed generally modest correlations, while OP exhibited anticorrelations. Finally, a subcortical-motor cluster, composed of the BG and MCE networks, showed moderate correlation with each other (r = 0.48) and weak associations with the visual-default cluster. Interestingly, their correlation patterns with other components were diametrically opposed: the BG network showed fair-to-moderate positive correlations with the multisensory cluster and negative correlation with SOM, while the MCE network displayed the reverse pattern.

### Human tICA

In the human dataset, tICA decomposition, 14 of the 20 components were categorized as functionally relevant based on their anatomical distribution and frequency spectra (Figure 5). These networks spanned primary sensory, association, and higher cognitive domains, enabling a direct comparison with the networks identified in marmosets.

**Figure 5.**
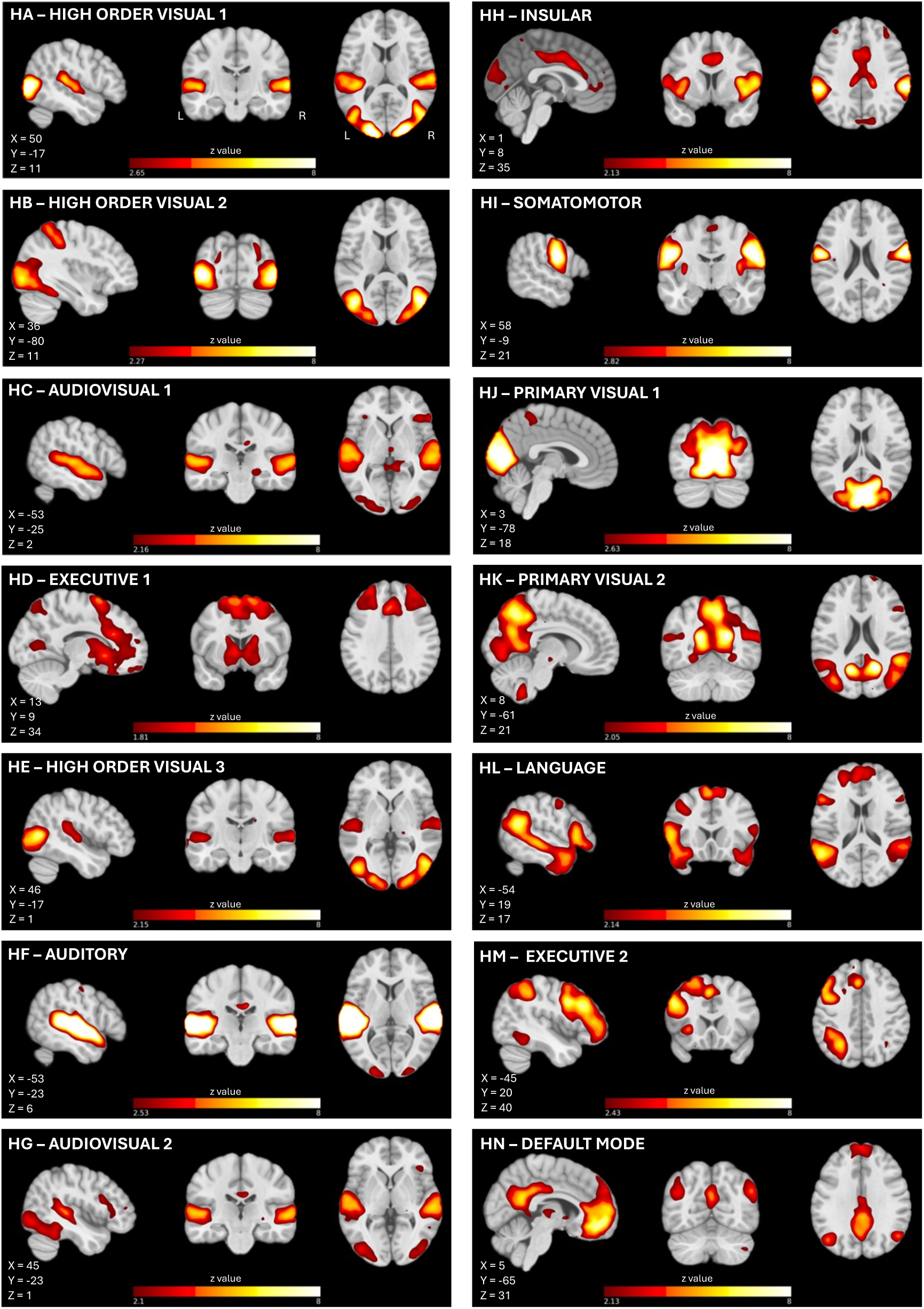
Spatial maps of the 14 functional human tICA components. Z-scored spatial maps of each human component are overlaid on the MNI152 template. MNI coordinates of the presented perspective are reported in the left bottom corner of each panel. Networks include higher-order visual (HOV), audiovisual (AV), auditory (AUD), executive (EXE), insular (INS), somatomotor (SOM), primary visual (PVIS), language (LAN), and default mode (DMN). Color indicates voxel-wise z-scores, thresholded at p < 0.5 (posterior probability that a voxel belongs to the active distribution rather than noise). Components were labeled based on spatial overlap with known functional networks and their correspondence to task content.

Three components—HA, HB, and HE—were grouped as **higher-order visual networks (HOV)**. These included bilateral activation in occipital and parietal cortices, consistent with dorsal and ventral visual streams. Notably, component HB engaged the frontal eye fields (FEF) and lateral intraparietal cortex, resembling a saccade or attentional control network.

Two components, HC and HG, were classified as **audiovisual networks (AV)**. These showed strong activation in the superior temporal sulcus (STS) and auditory cortex, and in the case of HC, also included prefrontal regions such as the inferior frontal gyrus (IFG), area 8Av, and PEF. Component HG extended into the retrosplenial complex (RSC), suggesting involvement in integrating auditory, visual, and contextual information.

Component HF was identified as a **pure auditory network (AUD)**, with strong activation in the auditory cortex, STS, and insula. Like HG, it overlapped with the RSC, reflecting potential convergence of sensory and default-related processes.

Two **executive networks (EXE)** were detected: HD, a bilateral network spanning dorsolateral prefrontal cortex, anterior cingulate, and parietal regions, also engaging the basal ganglia; and HM, a left-lateralized network including the dorsal stream, medial prefrontal cortex, and parietal cortex, potentially linked to the dorsal attention system (49–52).

The **insular network (INS)**, component HH, encompassed the anterior insula, anterior cingulate cortex, and PF complex—regions consistent with the salience network described in humans (53). **Somatomotor (SOM)** functions were represented by component HI, which showed strong bilateral activation in the precentral and postcentral gyri and aligned with typical somatomotor maps (49–52).

Early visual processing was captured by two **primary visual networks (PVIS)**—components HJ and HK—covering V1, V2, and adjacent areas. Notably, component HK has sometimes been considered part of the human default mode network due to its anatomical distribution, which includes the boundary between the medial occipital cortex and the posterior cingulate cortex (49,50).

Component HL, designated the **language network (LAN)**, activated frontal areas (left IFG, 44, 45, 47), superior frontal language area (SFL), posterior STS, and temporal pole areas (TPOJ, TG), with an additional cluster in the posterior cingulate (54,55). Its left-lateralized topology and co-activation of frontal-temporal language areas point to a role in speech comprehension and semantic processing during the movie.

Finally, component HN was identified as the **default mode network (DMN)**, encompassing the medial prefrontal cortex, PCC, angular gyrus, and hippocampus, consistent with canonical DMN architecture described in resting-state and naturalistic fMRI studies (49,50,56,57).

Voxel-wise probability maps (Figure 6A) indicated the most consistent recruitment across participants in auditory and visual cortices. By contrast, somatosensory and ventral temporal areas showed relatively sparse recruitment. These spatial patterns were not explained by signal quality, as confirmed by comparison with the tSNR map (Figure 6B), supporting the functional relevance of the extracted components.

**Figure 6.**
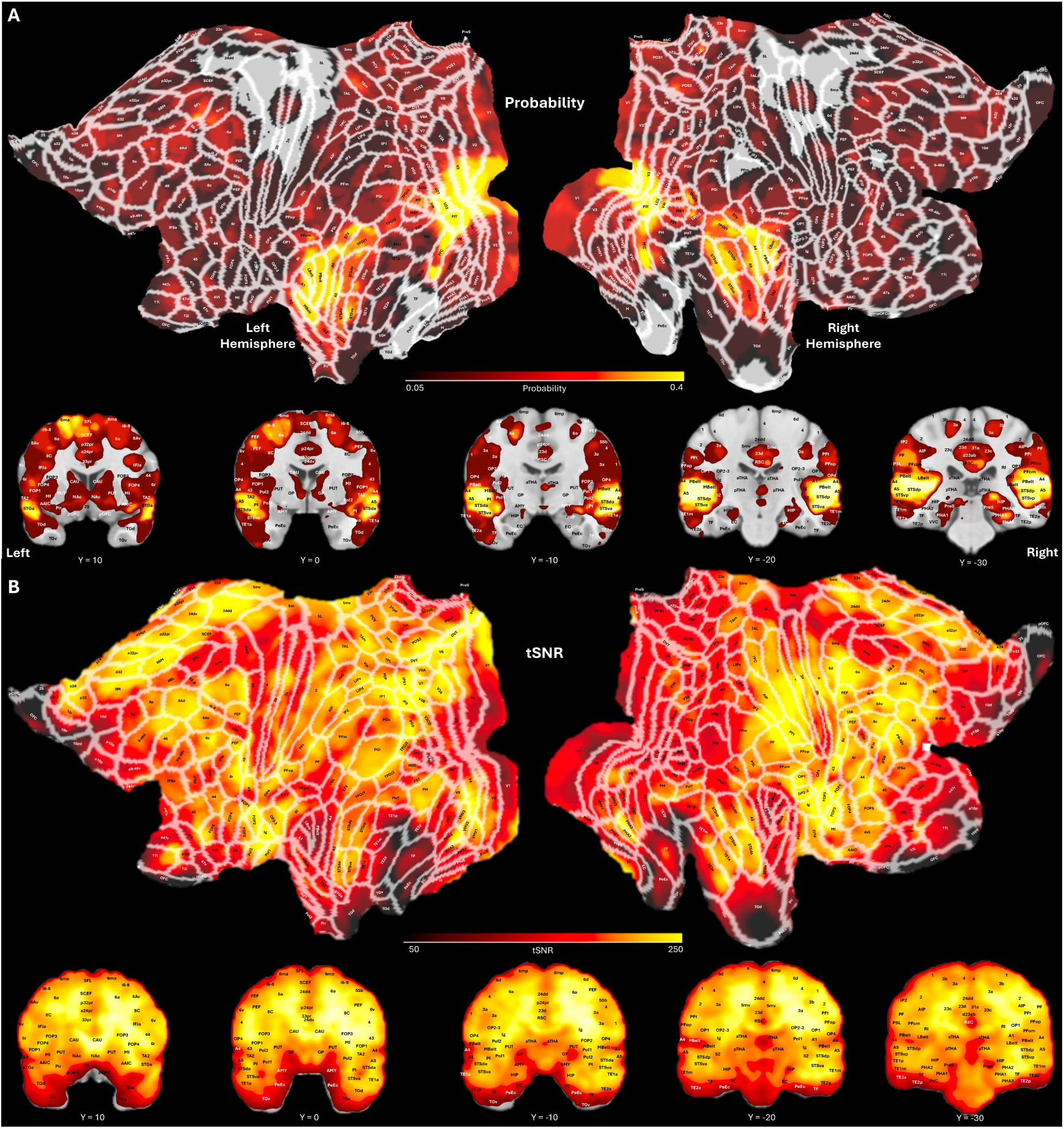
Spatial distribution and tSNR of human components. (A) Probability map showing the number of components in which each voxel was active across the 14 functional networks (thresholded as indicated in Figure 5). High-frequency recruitment was observed in auditory and visual areas, with sparse representation in somatosensory and ventral temporal cortex. (B) Mean tSNR map across all human participants. Visual inspection confirms that functional component distributions were not solely determined by tSNR patterns. Both maps are projected onto inflated and flat surface representations of the human brain, as well as onto coronal slices from an anatomical template in MNI space. White outlines indicate cortical areas based on the Human Connectome Project’s multi-modal parcellation atlas (77).

### Human Component Relationships

Hierarchical clustering of the 14 human functional components (Figure 7A) revealed three major groupings, reflected in the corresponding within-species correlation matrix (Figure 7B).

**Figure 7.**
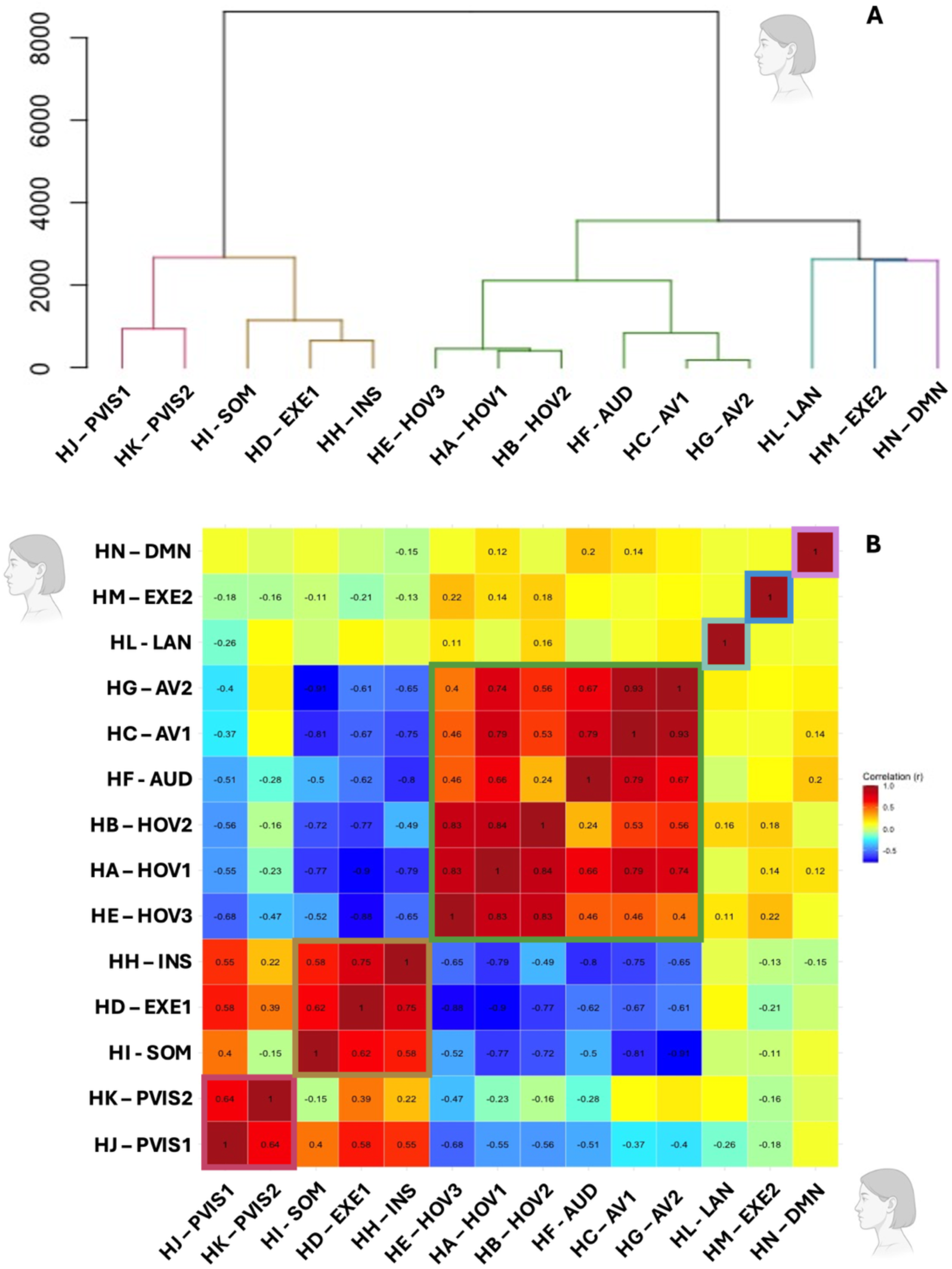
Human component clustering and temporal relationships. (A) Hierarchical clustering dendrogram of the 14 functional human components using squared Euclidean distance and Ward’s linkage. Clusters include primary visual (red), sensorimotor-executive-insular (gold), multisensory (green), and functionally distinct components (light blue, dark blue, purple). (B) Within-species correlation matrix showing pairwise temporal correlations (Pearson’s r) between component timecourses. Components are ordered based on the sequence specified by the hierarchical clustering tree. Sensorimotor and audiovisual networks show high internal correlation, while the language, executive-2, and DMN components are temporally distinct from other clusters. Clusters identified through hierarchical clustering are highlighted on the matrices with colored squares, following the same color scheme presented in panel A. Red shades indicate positive correlations, while blue shades represent negative correlations. Only correlations with an absolute r value greater than 0.10 are displayed.

The first branch of the dendrogram encompassed two functionally distinct but moderately linked clusters. The first comprised the two primary visual networks (PVIS1 and PVIS2), which were strongly correlated with each other (r = 0.64), reflecting their shared involvement in early visual processing. The second cluster was a **sensorimotor-executive-insular** group, comprising the somatomotor (SOM), executive 1 (EXE1), and insular (INS) networks, which also showed strong mutual correlations (r = 0.58–0.75), suggesting co-activation related to action observation, executive demands, or salience-driven processes. These two clusters were grouped together based on moderate cross-cluster correlations, most notably between SOM and PVIS1 (r = 0.40), though SOM was weakly anticorrelated with PVIS2 (r = –0.15), indicating some degree of functional separation.

The second branch was a **multisensory network group**, containing the HOV, AV, and AUD components. These networks were highly intercorrelated (r = 0.24–0.93), reflecting the integrated audiovisual nature of the stimulus and the simultaneous activation of visual and auditory pathways.

Three components—LAN, EXE2, and DMN—were more weakly correlated with other networks, forming functionally distinct entitie**s**. The LAN component showed minimal temporal overlap with either sensory or attentional networks, reflecting the specificity of its response to linguistic content. EXE2, the left-lateralized dorsal attention-like network, showed modest correlation with visual components and anticorrelation with EXE1, indicating a potentially distinct functional role. The DMN component exhibited little to no correlation with other networks, in line with its established role in internally oriented cognition.

### Cross-Species Correlations

To assess the degree of temporal alignment between functional brain networks across species, we computed the cross-species correlation matrix based on the timecourses of the 12 marmoset and 14 human functional components (Figure 8). This analysis revealed two major network groupings that differed in the strength and structure of their interspecies correspondence.

**Figure 8.**
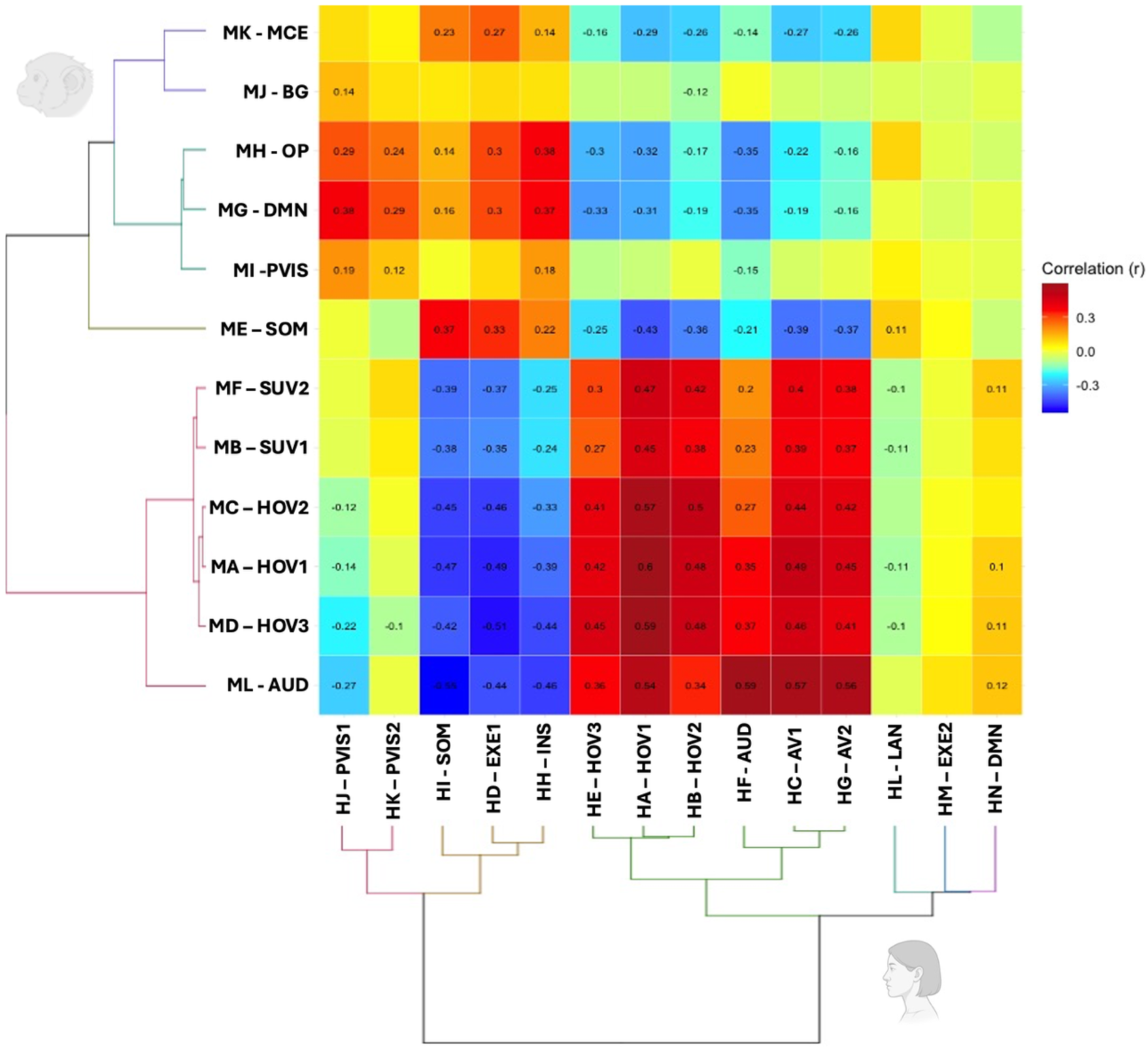
Cross-species correlation matrix. Matrix showing temporal correlations (Pearson’s r) between the 12 marmoset and 14 human functional components. Warm colors indicate positive correlation; cool colors indicate anticorrelation. Two major interspecies clusters emerge: a conserved multisensory cluster (upper left), including visual, auditory, and audiovisual networks, and a divergent cognitive cluster (lower right) comprising somatomotor, executive, and default mode components. Language and lateralized executive networks in humans show little or no correspondence in the marmoset. Only correlations with an absolute r value greater than 0.10 are displayed.

The first was a **conserved multisensory cluster**, comprising audiovisual and visual networks in both humans and marmosets. This group included the higher-order visual (HOV), subcortical-visual (SUV), auditory (AUD), and audiovisual (AV) components. Components within this cluster demonstrated modest to strong interspecies correlations, with values ranging from r = 0.27 to 0.60. Notably, the marmoset HOV and SUV components showed robust temporal alignment with human HOV and AV networks, highlighting convergent dynamics in visual processing streams. The marmoset AUD was most strongly correlated with the human AUD and AV networks, with cross-species correlations reaching r = 0.56–0.59. These results suggest that core sensory systems—particularly those involved in processing dynamic audiovisual input—exhibit shared temporal structure and functional organization across species when presented with naturalistic stimuli.

Despite substantial differences in cortical expansion and specialization, this cluster points to a high degree of evolutionary conservation in sensory integration. The inclusion of AV networks engaging multimodal integration regions, such as the STS in humans and pulvinar in marmosets, further suggests that the brain’s capacity to synchronize visual and auditory input under ecologically valid conditions may be a foundational property of primate neural systems.

The second major grouping represented a **divergent cognitive cluster**, encompassing components associated with somatomotor, executive, default mode, and subcortical-motor functions. This cluster showed only modest cross-species correlations (r = 0.12–0.38) and differed in composition between species. In humans, it included the somatomotor (SOM), insular (INS), and executive (EXE1) components, along with the PVIS network. In marmosets, the corresponding cluster included PVIS, SOM, OP, DMN, basal ganglia (BG), and motor-cerebellar (MCE) components. These findings suggest some shared temporal structure in sensorimotor and motor-associative processing, though the overall organization appeared more species-specific and variable in strength.

Importantly, several human components—particularly those associated with higher cognition—exhibited minimal or no correlation with any marmoset network. The **language network (LAN)**, characterized by strong left-lateralization and robust activation of frontal and temporal language areas, showed diffuse and weak correlations across all marmoset components. This was expected given the absence of linguistic capacity in non-human primates and the semantic specificity of the auditory content in the movie for human viewers.

Similarly, the **left-lateralized executive network (EXE2)** showed minimal alignment with any marmoset component and was weakly anticorrelated with the marmoset DMN and subcortical-motor clusters. This component likely reflects uniquely human attentional and goal-directed mechanisms, consistent with evolutionary expansions in lateral prefrontal cortex.

The **default mode network (DMN)** in humans also lacked a clear homolog in the marmoset data. Although a marmoset DMN component was identified (component MG), it did not correlate strongly with the human DMN and instead aligned more closely with marmoset visual and parietal networks and with the human primary visual, executive 1 and insular networks.

This suggests that while the anatomical topography of the DMN may be preserved across species, its functional dynamics and internal coherence during naturalistic viewing may differ significantly. This interpretation suggests that functional equivalence of the default mode network across primate species may be more limited than previously assumed.

Together, these findings reveal a dichotomy in cross-species brain dynamics. **Sensory networks show strong correspondence**, reflecting conserved processing of audiovisual input, while **higher-order cognitive networks diverge**, likely reflecting evolutionary changes in cortical organization and function. The ability of naturalistic fMRI paradigms to uncover these patterns demonstrates their utility in comparative neuroscience and reinforces the potential of the marmoset as a model for studying conserved sensory processes and the evolutionary emergence of complex cognition.

## Discussion

In this study, we used movie-driven functional magnetic resonance imaging (md-fMRI) to compare brain activity in humans and common marmosets as they watched the same naturalistic audiovisual stimulus. This paradigm provided a rich, ecologically valid context to assess cross-species functional network organization, expanding upon our earlier work identifying interspecies homologies in face-selective visual areas (14). By introducing diverse auditory and visual content — including conspecific and heterospecific vocalizations, human speech, and dynamic scenes — we engaged a broader range of sensory and associative regions, allowing for a more comprehensive examination of shared and divergent brain networks.

To analyze these responses, we applied tensor independent component analysis (tICA), a fully data-driven method that decomposes brain activity into temporally and spatially independent components without reliance on predefined network templates. This approach is well-suited to complex, continuous stimuli like movies, enabling the extraction of distinct functional networks that reflect moment-to-moment neural dynamics. Our analysis extracted 20 components per species, of which 12 in marmosets and 14 in humans were identified as functionally relevant (i.e., non-noise). We then compared temporal dynamics of these networks within and across species.

We observed strong cross-species correlations among core sensory systems, particularly those supporting audiovisual processing. Higher-order visual networks (HOV), subcortical-visual components (SUV), and auditory networks (AUD) formed tightly correlated clusters in both species, showing significant temporal alignment. These findings suggest a shared temporal structure in how primate brains process dynamic audiovisual input, despite differences in cortical architecture and specialization. The high degree of within- and between-species synchrony among these networks reinforces the idea that sensory integration, especially in naturalistic contexts, is supported by evolutionarily conserved mechanisms.

In contrast, components linked to higher cognitive functions — including language, executive control, and default mode processes — were uniquely observed in humans and did not correlate with any marmoset networks. The language network (LAN), for instance, exhibited strong left-lateralized recruitment of frontal and temporal regions traditionally associated with speech comprehension and semantic processing. Its absence in marmosets aligns with the lack of linguistic capacity in non-human primates and underscores the human specificity of this network. Similarly, a left-lateralized executive component resembling the dorsal attention network emerged only in humans, suggesting functional specialization in attentional control and top-down modulation not mirrored in the marmoset brain.

The internal organization of networks also differed markedly between species. While human components showed greater temporal segregation, with distinct correlation clusters representing sensory, cognitive, and default mode systems, marmoset components were more temporally synchronous. This pattern suggests a more distributed and overlapping functional organization in marmosets, where networks are less differentiated and more broadly engaged by naturalistic stimuli. Such differences may reflect evolutionary divergence in brain architecture and processing capacity, with the human brain exhibiting a higher degree of functional specialization.

Importantly, several findings confirmed and extended prior resting-state fMRI observations. In marmosets, components resembling default mode, somatomotor, visual, auditory, and basal ganglia networks closely aligned with those identified in earlier studies (12,13,45,46), lending support to the reproducibility and reliability of tICA under naturalistic stimulation. Moreover, the networks extracted recruit all the major functional hubs previously described in the awake marmoset brain (12,58). Although many of the major functional hubs identified in resting-state studies (e.g., caudate, putamen, thalamus, areas 8Av and TE3) are also highly recruited during movie viewing, others appear less involved in the networks extracted by tICA. This discrepancy raises the possibility that certain hubs play a more prominent role in organizing intrinsic brain activity during rest, but are less dynamically engaged during complex, multisensory stimulation. In other words, these areas may act as central nodes within the brain’s default communication architecture, but their functional influence may shift when the brain is engaged in externally driven, stimulus-bound processing. This distinction suggests that the role of functional hubs is not static, but rather dynamic and context-dependent, modulated by the cognitive and sensory demands of the environment.

In humans, canonical resting-state networks—including default mode, somatomotor, executive, and insular networks (12,49–52,56)—were robustly extracted, reinforcing their stability across task-free and naturalistic paradigms. However, the movie-driven context allowed for the emergence of additional components, particularly in sensory and multisensory domains, that are less frequently resolved in resting-state conditions. This highlights the utility of naturalistic stimuli for engaging diverse functional systems and enhancing component separability.

The dynamic nature of the movie facilitated the emergence of multisensory integration networks, especially in humans. Several audiovisual components centered on the superior temporal sulcus (STS), a region known for integrating visual and auditory input and implicated in speech and social perception (59). In contrast, marmosets showed fewer distinct audiovisual components, and these often involved subcortical regions such as the superior colliculus and pulvinar—structures known to contribute to multisensory processing in non-human primates (60,61). The presence of multisensory networks in both species underscores the shared computational demands imposed by naturalistic stimuli (30,31,62). Nevertheless, these differences suggests that cortical multisensory integration may be more anatomically and functionally elaborated in humans, possibly reflecting adaptations for processing complex communicative stimuli such as speech.

Despite the absence of direct somatosensory stimulation, both species exhibited well-defined somatomotor networks. These findings align with previous studies reporting strong intrinsic connectivity in these regions during rest (12,13,45,46,49,50,52) and support the notion that spontaneous fluctuations and action observation can activate sensorimotor systems. The content of the movie, which included scenes of movement, social interaction, and goal-directed behavior, may have engaged these networks via internal simulation mechanisms (63–65). This is consistent with evidence for action observation networks in both humans (66–68) and marmosets (26), suggesting that cross-species similarities in somatomotor engagement may reflect not only structural homology but shared computational roles during passive observation.

An intriguing aspect of our findings lies in the interpretation of the marmoset default mode network (DMN). While anatomically similar to the human DMN, the marmoset component labeled as DMN clustered with visual and occipitoparietal networks and showed strong temporal coupling with sensory and subcortical components—patterns not observed in humans. In humans, the DMN was largely functionally segregated, exhibiting weak correlations with sensory and executive systems. Furthermore, the marmoset DMN does not show temporal coupling with the human DMN, whereas it shows moderate correlations with the human executive and insular networks, which respectively resemble known cognitive control and salience networks in humans (49–51,56). This discrepancy highlights a critical gap in our understanding of the marmoset DMN: unlike the human DMN, which is extensively characterized in the literature (49,50,52,56,57), resting-state studies in marmosets report inconsistent findings, with some identifying networks that resemble the human DMN anatomically and others reporting divergent patterns (11–13,45–48).

These observations raise the possibility that the so-called marmoset DMN may serve a different function, perhaps related to visual attention or environmental monitoring, rather than internally directed thought or mind-wandering as in humans (69). The lack of interspecies correlation between DMN components, despite periods of reduced stimulation in the movie, underscores the need to reconsider the functional equivalence of these networks across species. Given the rich, naturalistic design and cross-species comparability of our paradigm, this work may offer a valuable framework to further clarify the architecture and role of the DMN in marmosets and its potential evolutionary divergence from the human counterpart.

Subcortical networks also revealed interesting cross-species differences. In marmosets, we identified a distinct basal ganglia (BG) component encompassing the caudate, putamen, and thalamus — structures frequently observed as independent networks in marmoset resting-state studies (12,13,45,46) and major functional hubs of the marmoset brain (58). This BG network showed strong correlations with subcortical-visual components and the primary visual network, suggesting integrated processing during movie viewing. In contrast, no basal ganglia component emerged from the human tICA decomposition. This discrepancy may reflect reduced detectability of subcortical signals in human imaging, or fundamental differences in the degree to which subcortical regions form temporally coherent networks. While some human studies do extract basal ganglia components (12,52,70,71), others do not (49–51), pointing to variability in detectability and functional coupling depending on species, analysis methods, and stimulus context.

A similar pattern was observed with cerebellar networks. The motor-cerebellar component in marmosets included motor cortex, cerebellum, and periaqueductal gray — regions seldom observed as a unified network in human resting-state studies. Its positive correlation with human somatomotor and cognitive control networks, and negative correlation with audiovisual systems, suggests a broader role in sensorimotor integration, potentially extending into cognitive domains. In humans, no cerebellar network was extracted, consistent with known difficulties in resolving posterior fossa activity in whole-brain ICA (49–52, but see 72). However, targeted ICA approaches have demonstrated the cerebellum’s rich functional architecture (73), and its absence here likely reflects limitations in sensitivity rather than true functional differences. Our findings suggest that naturalistic paradigms may facilitate the recovery of cerebellar activity, particularly in species like the marmoset where the cerebellum may play a more integrated role in behavior.

Finally, in addition to the previously discussed default mode network, our analysis revealed two other human networks—the language and executive 2 components—that lacked clear analogues in the marmoset dataset. Their absence could reflect several factors: species differences in anatomical expansion and specialization; divergence in cognitive capacity; and differences in the relevance of the movie content. For instance, the language network (74) likely emerged due to the presence of continuous speech, which holds semantic value for human viewers but likely lacks meaningful content for marmosets. The executive component, strongly lateralized and resembling the dorsal attention stream (49–52,75), may reflect uniquely human mechanisms for sustained attention and goal-directed processing. The emergence of such networks highlights the sensitivity of md-fMRI and tICA to cognitive specialization and underscores the value of cross-species comparison in understanding the evolution of brain function.

In summary, our findings demonstrate that movie-driven fMRI, when combined with data-driven analytical techniques like tICA, provides a powerful platform for mapping functional brain organization across species. This approach enables the identification of both conserved and divergent neural dynamics, offering insight into the evolutionary underpinnings of sensory integration, cognitive specialization, and large-scale brain network architecture. By making our data and tools freely available, we aim to promote broader use of the marmoset as a translational model and foster new avenues for comparative neuroscience.

### Limitations

While our current movie-driven fMRI approach offers a rich and ecologically valid stimulus, it may nevertheless be too complex for precise functional localization. However, more targeted movies could be designed to emphasize specific features, such as increased frequency of conspecific calls or varying levels of motion and contrast. It is also possible to focus analyses on selected segments of the movie to isolate responses to particular stimulus categories (e.g., faces, bodies, or vocalizations).

Another limitation is that brain regions with closely related functions—such as those along the marmoset’s occipito-temporal axis—may be grouped together due to high temporal overlap in their activity patterns. In cases where finer functional distinctions are required, more targeted approaches such as functional localizers, electrophysiological recordings, or the use of more specifically designed movie stimuli may be necessary.

## Acknowledgements

Support was provided by a Discovery grant by the Natural Sciences and Engineering Research Council of Canada and the Canadian Institutes of Health Research (FRN 183973). We also acknowledge the support of the Government of Canada’s New Frontiers in Research Fund (NFRF), [NFRF-T-2022-00051]. We wish to thank Cheryl Vander Tuin, Whitney Froese, Hannah Pettypiece, and Miranda Bellyou for animal preparation and care, Dr. Alex Li and Trevor Szekeres for scanning assistance, Dr. Kyle Gilbert, and Peter Zeman for coil designs.

## Data Availability

The individual maps for the human and marmoset samples, along with the code underpinning this study, are fully available online on Zenodo: https://doi.org/10.5281/zenodo.12746414.

